# Single-cell imaging reveals that *Staphylococcus aureus* is highly competitive against *Pseudomonas aeruginosa* on surfaces

**DOI:** 10.1101/2021.06.28.450150

**Authors:** Selina Niggli, Tobias Wechsler, Rolf Kümmerli

## Abstract

*Pseudomonas aeruginosa* and *Staphylococcus aureus* frequently occur together in polymicrobial infections, and their interactions can complicate disease progression as well as treatment options. While interactions between *P. aeruginosa* and *S. aureus* have been extensively described using planktonic batch cultures, little is known about whether and how individual cells interact with each other on solid substrates. This is important, because in infections, both species frequently colonize surfaces to form microcolony aggregates and biofilms. Here, we performed single-cell time-lapse fluorescence microscopy combined with automated image analysis to describe interactions between *P. aeruginosa* PAO1 with three different *S. aureus* strains (Cowan I, 6850, JE2) during microcolony growth on agarose surfaces. While *P. aeruginosa* is usually considered the dominant species, we found that the competitive balance tips in favor of *S. aureus* on surfaces. We observed that all *S. aureus* strains accelerated the onset of microcolony growth in competition with *P. aeruginosa* and significantly compromised *P. aeruginosa* growth prior to physical contact. These results suggest that *S. aureus* deploys mechanisms of both resource competition and interference competition via diffusible compounds. JE2 was the most competitive *S. aureus* strain, simply usurping *P. aeruginosa* microcolonies when coming into direct contact, while 6850 was the weakest competitor itself suppressed by *P. aeruginosa*. Moreover, *P. aeruginosa* reacted to the assault of *S. aureus* by showing increased directional growth and expedited expression of quorum sensing regulators controlling the synthesis of competitive traits. Altogether, our results reveal that quantitative single-cell live imaging has the potential to uncover microbial behaviors that cannot be predicted from batch culture studies, and thereby contribute to our understanding of interactions between pathogens that co-colonize host-associated surfaces during polymicrobial infections.

## Introduction

Bacterial infections are frequently caused by multiple species, and such polymicrobial infections can be more virulent and more difficult to treat (1, 2). For this reason, there is great interest in understanding how pathogens interact and how their interactions affect virulence and treatment outcomes (3, 4). *Pseudomonas aeruginosa* (PA) and *Staphylococcus aureus* (SA) have emerged as a particularly important model system in this context (5–7), as these two pathogens co-occur in multiple types of infections, including the lungs of cystic fibrosis (CF) patients and chronic wounds (8–11).

Interactions between PA and SA have been studied at the molecular, ecological, and evolutionary levels. Molecular studies revealed that interactions between PA and SA seem to be predominantly antagonistic, whereby PA is the dominant species suppressing the growth of SA through the production of a variety of inhibitory molecules like proteases, biosurfactants, siderophores, and toxic compounds (12–19). At the ecological level, it was shown that the strain genetic background, the spatial structure of the environment, and the relative frequency of strains impact the outcome of interactions (20–22). For example, in our previous work, we showed that PA can only displace the SA strain JE2 when occurring above a certain threshold frequency but fails to invade SA populations when being initially rare (below 5%). At the evolutionary level, there is great interest to understand whether PA and SA (co-)evolve (23–26) and indeed, there is evidence that this is the case, with PA becoming either more (27) or less (24, 28–30) competitive towards SA over time. Important to note is that our understanding of PA and SA interactions is predominantly based on laboratory batch culture experiments, where large populations grow under shaken conditions. This contrasts with the environment prevailing in infections, where PA and SA frequently act as surface-colonizing pathogens, forming small microcolony aggregates and biofilms (31–37). It is conceivable to assume that interspecies interactions mainly take place at the front of such bacterial aggregates, and that interactions therefore occur at the local micrometer, and not the batch culture scale. Yet little is known about the dynamics and the outcome of competition between species at this scale. The single-cell study by (38) is a notable exception, where it was shown that PA modifies its motility upon sensing nearby SA cells.

In our study, we aim to deepen our understanding of single-cell interactions between PA and SA both at the behavioral and fitness level. For this purpose, we performed time-lapse fluorescence microscopy, where we tracked growing microcolonies on solid agarose patches, either in mono- or mixed culture. Using automated image analysis, we quantified the time until the onset of growth of a microcolony and the number of progenies produced per founder cell, and tested whether these two fitness metrics were influenced by the presence of a competitor. Next, we assessed whether there is growth directionality in mixed cultures, whereby the competing species would grow towards or away from each other. We then followed physical encounters between microcolonies of the two species and allocated the various interaction patterns observed into distinct behavioral categories. In a final experiment, we focused on PA and asked whether PA reacts to the presence of SA by changing the expression of key quorum sensing (QS, cell-to-cell communication) genes, known to regulate competitive traits against SA (5, 39). Important to note is that we repeated all experiments for three SA strains (Cowan I, 6850, JE2) in competition against a single PA strain (PAO1) to test whether micro-scale interactions are SA strain- specific.

## Materials and Methods

### Bacterial strains used and general growth conditions

We used tagged variants of the *Pseudomonas aeruginosa* (PA) strain PAO1 and the untagged *Staphylococcus aureus* (SA) strains Cowan I, 6850 and JE2 (also outlined in Table S1) for all experiments. For time-lapse experiments, we used a constitutively expressed green fluorescent protein (*attTn7::ptac-gfp*) in the chromosome of PA as a marker to distinguish PA from SA. For experiments with PA gene reporters, we used PA strains carrying constructs with promoters of interest fused to *mCherry* together with the housekeeping gene promoter of *rpsL* fused to *gfp* (*attTn7::lasR-mCherry;rpsL-gfp* and *attTn7::rhlR-mCherry;rpsL-gfp*) (40). Prior to imaging, bacterial overnight cultures were grown in 10 ml tryptic soy broth (TSB, Becton Dickinson) in 50 ml falcon tubes for ± 16 hours at 37 °C and 220 rpm with aeration. After centrifugation and removal of the supernatant, we washed bacterial cells using 10 ml 0.8% NaCl solution and adjusted OD_600_ (optical density at 600 nm) to obtain similar cell numbers per ml for both PA and SA. This was achieved by adjusting OD_600_ of PA to 0.35, for SA strains JE2 and 6850 to 0.65 and for Cowan I to 0.85. Samples were diluted 1:10 with 0.8% NaCl and PA-SA strain pair combinations were mixed at a ratio of 1:1. 1.5 µl of this mix and of the respective monocultures was used to inoculate agarose pads for microscopy.

**Table 1.** Summary of effects in mixed microcolonies compared to monoculture microcolonies.

|  | Growth onset | Overall growth | Directionality of growth | Behavior when mixed colonies come into close contact | *lasR* gene expression | *rhIR* gene expression |
| --- | --- | --- | --- | --- | --- | --- |
| PA + Cowan I | Delayed*** | Reduced*** | * | 35% SA overgrows PA, 65% PA grows around SA | Timepoint 1 increased, Timepoint 2 decreased | Timepoint 1 increased, Timepoint 2 decreased |
|  | Accelerated** | n.s. | n.s. |  |  |  |
| PA + 6850 | n.s. | Reduced*** | * | 18% SA overgrows PA, 82% PA grows around SA | Timepoints 1 and 2 increased | Timepoints 1 and 2 increased |
|  | Accelerated** | Reduced*** | n.s. |  |  |  |
| PA + JE2 | n.s. | Reduced*** | * | 92% SA overgrows PA, 8% PA grows around SA | Timepoint 1 increased, Timepoint 2 decreased | Timepoints 1 and 2 increased |
|  | Accelerated** | n.s. | n.s. |  |  |  |

### Preparation of microscope slides for imaging

The following method was previously described and successfully used in our laboratory (41). To prepare agarose pads, we used standard microscopy slides (76 mm x 26 mm), standard coverslips and ‘gene frames’ (Thermo Fisher Scientific). Each frame is 0.25 mm thick and sticky on both sides. As a solid growth substrate for bacteria, we heated 20 ml of TSB + 1% agarose in a microwave and pipetted an excess (ca. 400 µl) of medium into the gene frame chamber. We covered the chamber with a microscope coverslip and let the TSB + 1% agarose solidify for around 20 min. at room temperature. After solidification, we removed the coverslip by carefully sliding it upwards and divided the agarose pad into four smaller pads using a sterile scalpel. Channels were introduced around each pad to allow continuous supply of oxygen during microcolony growth. Finally, we pipetted 1.5 µl PA monoculture, 1.5 µl SA monoculture and two times 1.5 µl mixed culture on the four smaller pads. After evaporation of the droplet containing bacteria on the pad (ca. 3 min.), we sealed the pads with a new coverslip. Imaging or incubation of the agarose pads at 37 °C was started right after slide preparation was completed.

### Microcolony imaging in time-lapse and individual timepoint experiments

All microscopy experiments were carried out at the Center for Microscopy and Image Analysis of the University of Zurich (ZMB) with a widefield Olympus ScanR HCS system and the Olympus cellSens software. This microscope features a motorized Z-drive, a Lumencor SpectraX light engine LED illumination system and a Hamamatsu ORCA-FLASH 4.0 V2 camera system (16-bit depth and 2048 x 2048 resolution). For all experiments, we used a PLAPON 60x phase oil objective (NA = 1.42, WD = 0.15 mm) with double digital magnification.

For time-lapse microscopy, we imaged growing microcolonies with phase contrast (exposure time 100 ms) and FITC SEM (exposure time 50 ms, excitation = BP 470 ± 24 nm, emission = BP 515 ± 30 nm and DM = 485). Time-lapse recording was performed with temperature in the incubation chamber set to 37 °C for six hours with images taken every 10 min. We imaged one PA-SA strain combination per time-lapse experiment and repeated this on three separate days (resulting in nine experiments). On each day, we imaged at least one field of view per monoculture and at least three fields of view for co-cultures.

For imaging individual timepoints to measure gene expression with the PA gene double reporters in the presence vs. absence of SA (5 hours and 8 hours after preparation and incubation of agarose pads at 37 °C), we used phase contrast (exposure time 100 ms), FITC SEM (exposure time 50 ms, excitation = BP 470 ± 24 nm, emission = BP 515 ± 30 nm and DM = 485) and TRITC SEM (exposure time 400 ms, excitation = BP 550 ± 15 nm, emission BP 595 ± 40 nm and DM = 558). We imaged both PA gene double reporters together with and without the three SA strains and the untagged controls (with and without SA) in three independent experiments. For every timepoint, we imaged four fields of view per strain combination and four blank positions (with no bacteria present) to use for average blank subtraction during image analysis (see below).

### Image analysis and quantification of growth and behavioral patterns

In a first step, we drift-corrected our time-lapse images in Fiji (42) using a drift correction script, published under a GNU general public license (https://github.com/fiji/Correct_3D_Drift). The drift-corrected images were then cropped to remove black space that was created during drift correction. Next, we exported the time-lapse series with the ilastik Import Export plugin as HDF5 (https://github.com/ilastik/ilastik4ij/). In ilastik (version 1.3.2), we created a pixel classification and object classification project in which we imported the respective HDF5 files (43). We segmented cells based on phase contrast (to distinguish cells from background) and gfp (to distinguish gfp-positive PA from gfp-negative SA cells), created the respective object predictions using a Gaussian blur with a sigma value of 0.5, the simple thresholding method with a threshold of 0.5, excluded objects smaller than 50 pixels, and exported the resulting object information. The remaining steps of our image analysis workflow were performed in R studio (version 3.6.3). First, we loaded the object predictions into a Shiny app that was programmed in our laboratory. This app allowed us to perform several steps. (1) Mark and exclude false positive cells; (2) exclude cells that exit or enter the field of view during imaging; (3) define groups of cells based on a hierarchical cluster analysis of the euclidean distance between the cells, which can be manually modified and corrected after visual inspection if necessary; and (4) calculate the center of mass for each cell group at each timestep using the formula x_COM = sum(x_i_ × a_i_) / sum(a_i_), where x_COM is the center of mass x, x_i_ is the cell position and a_i_ is the area of the cell. We used the information obtained from the Shiny app to calculate: (a) The onset of cell division for each microcolony; (b) the number of progeny cells per founder cell of a microcolony; and (c) the directionality of microcolony growth over time (see detailed descriptions below).

(a) To quantify the onset of cell division for each microcolony, we used the initial cell number of a group (N_i_) and calculated at which timestep of imaging that number exceeded N_i_ for the first time. (b) To calculate the number of progenies per founder cell N_p_, we used the formula N_p_ = (N_f_ – N_i_) / N_i_, where N_f_ is the final and N_i_ is the initial cell number of a group, respectively. (c) To calculate growth directionality D_g_, we used the formula D_g_ = D_e_ / D_a_, where D_e_ is the euclidean distance (corresponding to the distance between the two center of masses of a colony in the first and the last frame) and where D_a_ is the accumulated distance (corresponding to the sum of distances between the center of masses of a colony across all successive time points imaged). Random colony movements would lead to large D_a_ but low D_e_ distances, and thus lead to low directionality D_g_ values. In contrast, D_g_ values close to 1.0 would indicate high directional movement of a colony. The same formula to calculate growth directionality has previously been used by (38).

To quantify the different behavioral growth patterns in mixed microcolonies when PA and SA came into close contact, we manually screened all the time-lapse series and counted the two most distinct events: (1) SA pushes aside and overgrows PA and (2) PA grows around SA until the end of the six hours imaging period.

### Image processing to quantify PA gene expression

We segmented PA cells based on their constitutive gfp fluorescence (SA cells are non-fluorescent) using the interactive pixel and object classification workflow in ilastik (version 1.3.2) (43). We again applied a Gaussian blur with a sigma value of 0.5 and used the simple thresholding method with a threshold of 0.5. The resulting binary images (as png-files) were then exported and used as masks for mCherry and gfp quantification in Fiji. To do so, we used a custom-built script that uses the object predictions (in the form of binary images) created in ilastik, the blank images (to subtract average blank fluorescence), FITC (gfp) and TRITC (mCherry) channel images to quantify fluorescence in all predicted objects (corresponding to PA cells). This script performs the following steps: (1) Average blank subtraction in FITC and TRITC channels for each image (to correct for intensity differences across the field of view caused by microscope vignetting) ; (2) image cropping to a region of interest where all cells are well focused; and (3) background subtraction for both fluorescent channels in each cropped image (to correct for background autofluorescence). We imported the resulting information about the objects (corresponding to PA cells) into R studio and then performed the following steps. (1) Removing objects with an area smaller than 0.5 µm^2^ (which are most likely not cells); (2) Adding a value of 1.0 to each integrated density value to make all datapoints positive (the integrated density is the mean grey value, corresponding to gfp or mCherry fluorescence, multiplied by the area of the cell); (3) calculating the log_10_ of all integrated density values; (4) subtracting autofluorescence of PA gene reporter strains growing alone and PA gene reporter strains growing together with SA (using the average log_10_ fluorescence intensity of the untagged PA strain growing alone and the untagged PA strain growing together with SA from the same timepoint and experiment, respectively); and (5) plotting the ‘corrected’ log_10_ integrated density for mcherry (TRITC) and gfp (FITC).

### Statistical analysis

All statistical analyses were performed with R Studio (version 3.6.3). To test whether SA influences PA growth (onset of growth, number of progenies per founder cell, growth directionality), we used analysis of co-variance (ANCOVA), where we fitted the culture type (PA alone, PA + Cowan I, PA + 6850, PA + JE2) as a fixed factor and the total number of microcolonies present in a field of view as a covariate. To test whether PA influences SA growth, we also used ANCOVA, but fitted SA strain genetic background (Cowan I, 6850, JE2) and presence/absence of PA as fixed factors and the total number of microcolonies present in a field of view as a covariate. Note that we log-transformed the response variable ‘number of progenies per founder cell’ to obtain normally distributed residues for statistical analysis.

To compare PA gene expression patterns across culturing conditions, we fitted the culture type (PA alone, PA + Cowan I, PA + 6850, PA + JE2) and the timepoint as fixed factors and ‘experimental block’ as additional factor (without interaction) to account for variation between independent experiments. The response variables (gfp and mCherry fluorescence values) were log_10_-transformed prior to statistical analysis (see above). For all data sets, we consulted diagnostic Q-Q plots and results from the Shapiro-Wilk test prior to statistical analysis to ensure that model residues are normally distributed. P-values were corrected using the false discovery rate method whenever necessary.

We used Fisher’s exact test to compare whether frequencies of behavioral patterns between PA and SA differ among SA strain background (Cowan I, 6850, JE2).

## Results

### *P. aeruginosa* fitness is compromised by *S. aureus* in a strain-specific manner

To address whether fitness of the two species is affected when growing together on a solid surface, we performed single-cell time-lapse microscopy of *P. aeruginosa* PAO1 (PA) alone or in the presence of the *S. aureus* (SA) strains Cowan I, 6850 or JE2 (Figure 1 and supplementary table 1). As proxies for fitness, we calculated (i) the onset of microcolony growth (i.e. time to first cell division) and (ii) overall microcolony growth (i.e. number of progenies per founder cell).

**Figure 1.**
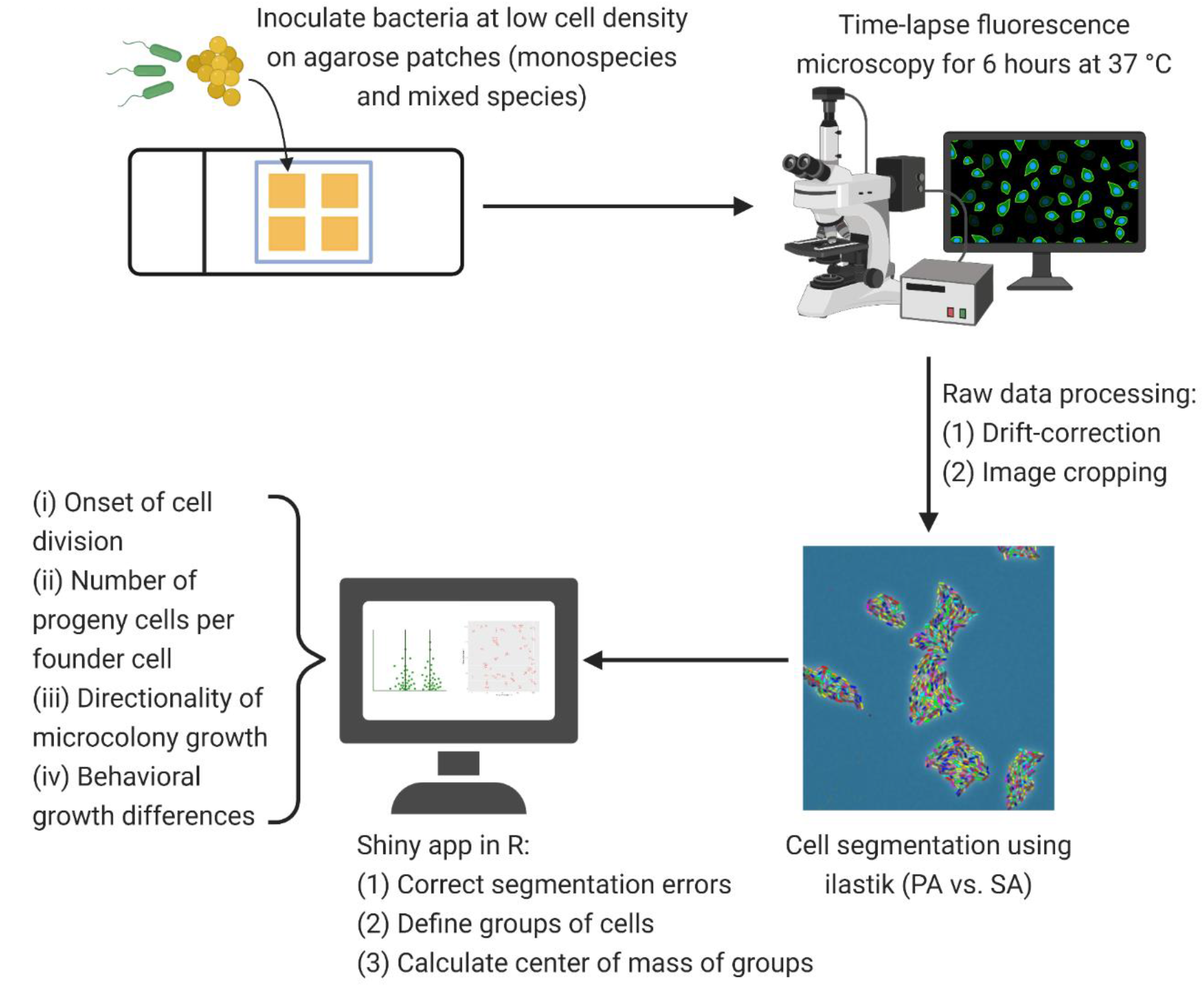
Microscopy workflow for time-lapse fluorescence microscopy. After adjusting *P. aeruginosa* (PA) and *S. aureus* (SA) to similar cell numbers, we inoculated bacteria (each species alone or mixed 1:1) at low cell density on TSB + 1% agarose patches. Time-lapse fluorescence microscopy was carried out for six hours at 37 °C with pictures taken every ten minutes. We drift-corrected and cropped the time-lapse images before cell segmentation (PA vs. SA) in ilastik (version 1.3.2). Using a Shiny app in R, we corrected segmentation errors from the exported object predictions, defined groups of cells and extracted the center of mass per group of cells for each timepoint. We then calculated the onset of growth (time to first cell division per microcolony), the number of progenies per founder cell in a microcolony, and the directionality of microcolony growth using automated scripts in R. Distinct microcolony interaction patterns between the species were manually assessed and counted.

We found that the onset of PA microcolony growth was significantly delayed in the presence of Cowan I (t_347_ = 6.42, p < 0.0001), but was neither affected by 6850 (t_347_ = 0.59, p = 0.5564) nor by JE2 (t_347_ = 1.90, p = 0.0875) (Figure 2a). In contrast, the presence of all three SA strains significantly reduced the number of progenies per founder cell for PA microcolonies (ANOVA: F_3,347_ = 8.58, p < 0.0001, Figure 2c). Interestingly, there were opposing effects of the number of microcolonies (sum of PA and SA microcolonies) in the field of view on the onset of growth and number of PA progenies. While higher numbers of microcolonies led PA to start dividing earlier, the number of PA progenies was reduced (ANOVA for the onset of growth: F_1,347_ = 6.94, p = 0.0088; number of progenies: F_1,347_ = 24.88, p < 0.0001). Overall, these findings show that PA fitness is compromised by the presence of SA in a strain-specific and cell density-dependent manner.

**Figure 2.**
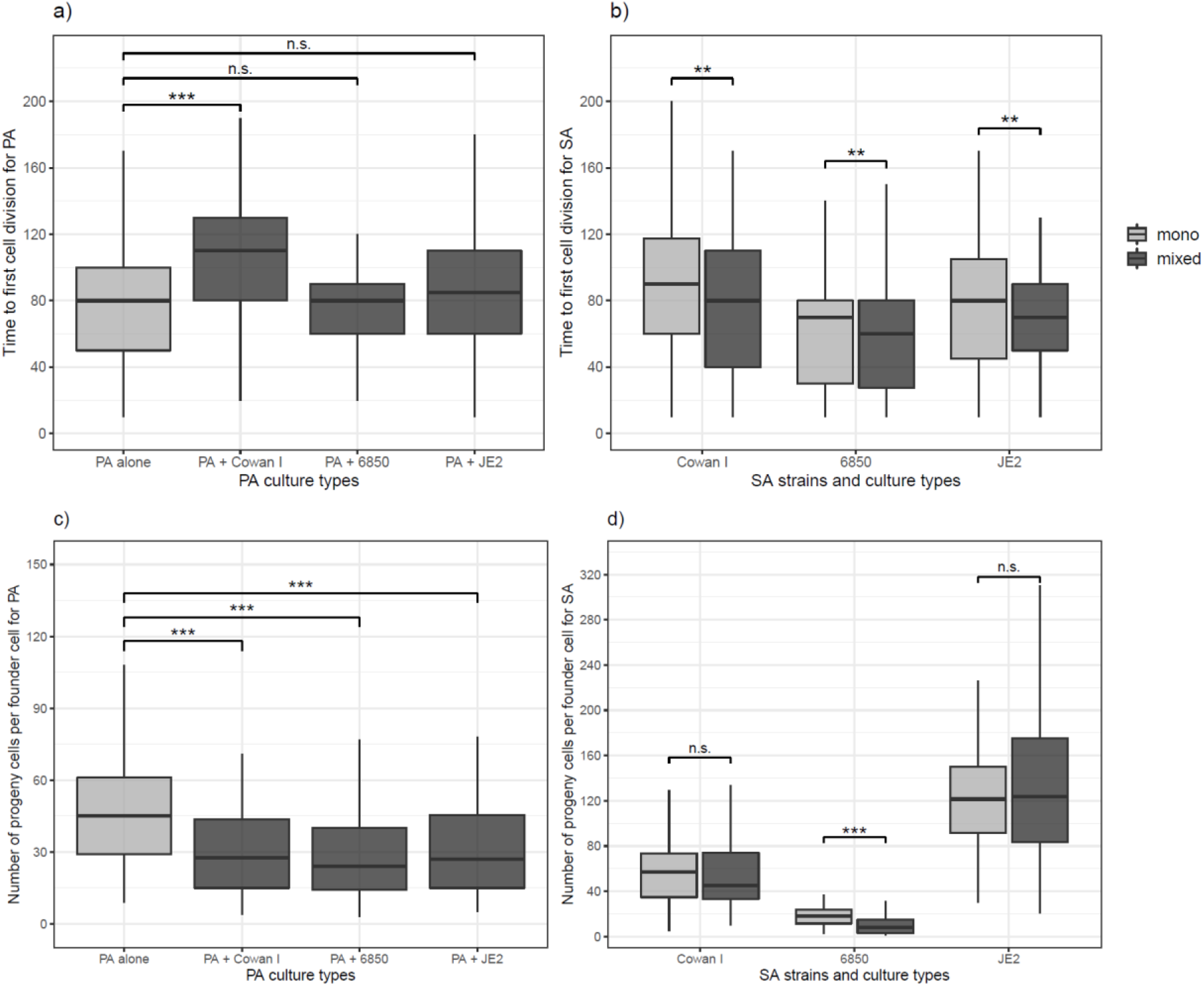
Time of first cell division and number of progenies per founder cell for *P. aeruginosa* (PA) and *S. aureus* (SA) microcolonies growing in mono- (light-grey) and mixed culture (dark-grey). a) Onset of cell division in PA microcolonies is significantly delayed in the presence of Cowan I, but not affected in the presence of 6850 and JE2. b) Onset of cell division in SA microcolonies is significantly accelerated in the presence of PA for all three SA strains. c) Number of PA progenies is significantly reduced in the presence of all three SA strains. d) Number of SA progenies is reduced in the presence of PA only for 6850, while the growth of Cowan I and JE2 remained unaffected. The box plots show the median (bold line) with the first and the third quartiles. The whiskers cover the 1.5* inter-quartile range (IQR) or extend from the lowest to the highest value if they fall within the 1.5* IQR. *** p < 0.001, ** p < 0.01, n.s., not significant. Data is from three independent experiments per PA-SA combination, with a total of 352 and 323 microcolonies for PA and SA strains, respectively.

### The onset of *S. aureus* microcolony growth is accelerated in the presence of *P. aeruginosa*

Next, we analyzed the fitness of SA strains from the same microscopy co-culture experiments as above. We found that the onset of microcolony growth depended on the SA strain (ANOVA: F_2,318_ = 8.09, p = 0.0004) and on the presence vs. absence of PA (F_1,318_ = 6.96, p = 0.0087). Particularly, we found that the presence of PA boosted the onset of SA microcolony growth (Figure 2b), while a higher number of microcolonies present in a field of view delayed it (F_1,318_ = 5.89, p = 0.0158). Comparisons of the number of progenies produced yielded significant differences between SA strains (F_2,317_ = 354.98, p < 0.0001) and a significant interaction between SA strain background and the presence vs. absence of PA (F_2,317_ = 13.35, p < 0.0001). The interaction is explained by the fact that the presence of PA reduced the number of progenies of 6850 (t_317_ = -4.27, p < 0.0001) but not of Cowan I (t_317_ = -0.36, p = 0.7180) or JE2 (t_317_ = 0.66, p = 0.7180, Figure 2d). Overall, these results show that the presence of PA accelerates the onset of SA growth on surfaces, whereas overall microcolony growth was either not affected or reduced (for 6850).

### *P. aeruginosa* but not *S. aureus* shows directional growth in the presence of a competitor

We further explored whether PA and SA show increased directional microcolony growth (away or towards each other) in the presence of a competitor. PA generally showed higher levels of directional growth than SA (Figure 3a), but directionality was only marginally increased in the presence of SA strains (ANOVA: F_3,348_ = 2.46, p = 0.0628). When repeating the analysis with a simpler statistical model testing whether PA shows directional growth in the presence of SA overall (i.e. collapsing SA factor levels), we found indeed significantly increased directional growth (F_1,350_ = 4.02, p = 0.0458), but the effect size was relatively small (mean directionality ± standard error of PA alone vs. with SA: 0.32 ± 0.02 vs. 0.36 ± 0.01).

**Figure 3.**
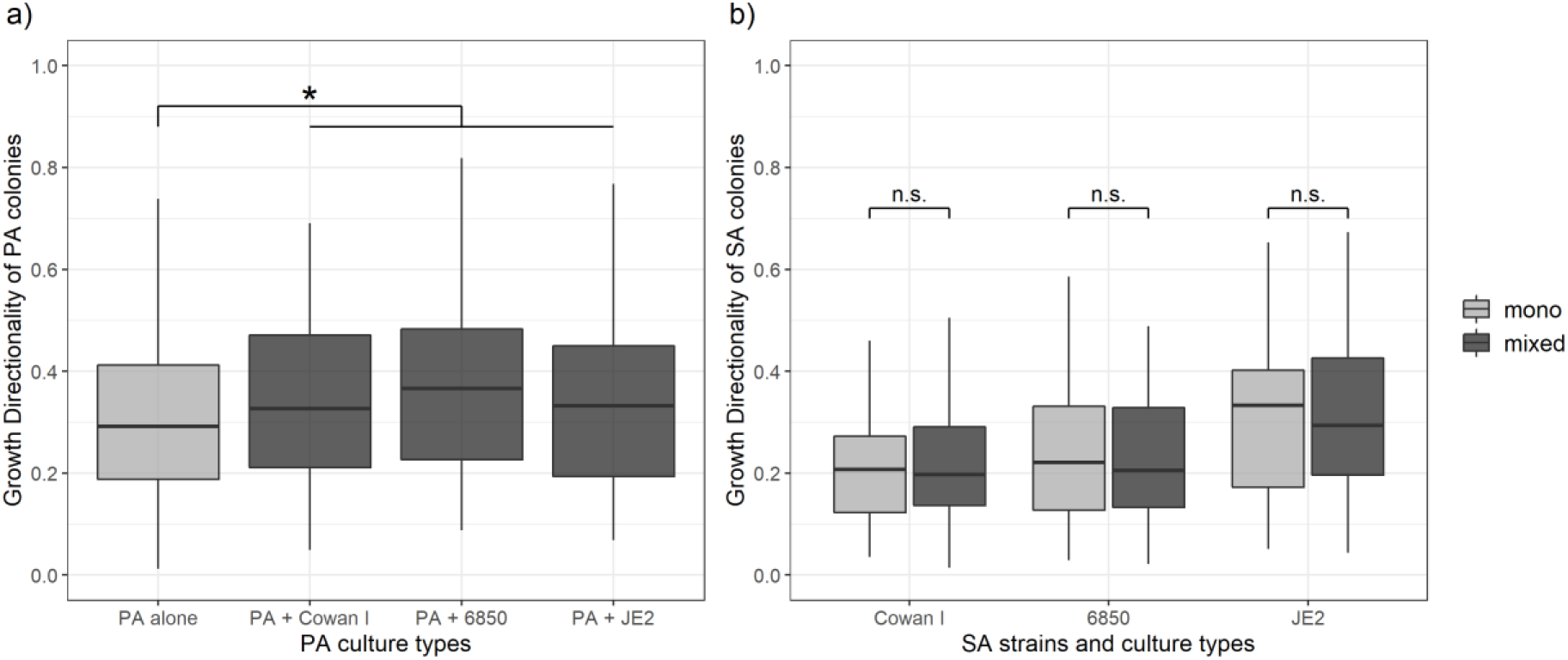
Directionality of microcolony growth of *P. aeruginosa* (PA) and *S. aureus* (SA) in monoculture (light-grey) and mixed culture (dark-grey). We calculated the directionality of growth as D_e_/D_a_ (where D_e_ is the euclidean and D_a_ is the accumulated distance, respectively). The closer this ratio is to 1.0, the more directional the movement of a microcolony is. a) Growth directionality of PA microcolonies is significantly increased in the presence of SA. b) Growth directionality of SA microcolonies is not affected by the presence of PA for none of the three SA strains. The box plots show the median (bold line) with the first and the third quartiles. The whiskers cover the 1.5* inter-quartile range (IQR) or extend from the lowest to the highest value if they fall within the 1.5* IQR. * p < 0.05, n.s., not significant. Data is from three independent experiments per PA-SA combination, with a total of 352 and 323 microcolonies for PA and SA strains, respectively.

For SA, directionality of growth was strain-dependent (ANOVA: F_2,319_ = 9.37, p = 0.0001), with JE2 growing more directional than Cowan I and 6850 (JE2 vs. Cowan I: t_319_ = 4.04, p = 0.0002; JE2 vs. 6850: t_319_ = 3.24, p = 0.0020; Cowan I vs. 6850: t_319_ = 0.44, p = 0.6620). Growth directionality was not affected by the presence of PA (ANOVA: F_1,319_ = 0.03, p = 0.8667, Figure 3b). Note that the number of microcolonies present per field of view did not have a significant effect on SA and PA growth directionality, and this covariate was thus removed from the statistical models. Overall, our analyses revealed that PA shows a weak but significant increase in directional growth in the presence of SA, whereas SA does not.

### Strain-specific interactions upon physical contact between microcolonies

By manually screening all the time-lapse images of our experiments, we noted two frequent behavioral interaction types upon physical contact between PA and SA (Figure 4): (1) PA grows around SA microcolonies, which can result in a ring-like structure that remains until the termination of imaging after six hours (movie S2 and S3) and; (2) PA comes in touch with SA, which results in PA growth arrest followed by SA pushing PA cells aside and (sometimes) overgrowing them completely (movie S4). We detected 62 distinctive instances in which PA either grows around SA (scenario 1: 29 cases, 46.8%) or is pushed aside and overgrown by SA (scenario 2: 33 cases, 53.2%). The frequency of these two PA behavioral patterns significantly differed in interaction with the three different SA strains (Fisher’s exact test p < 0.0001). While in the majority of cases, PA grew around microcolonies of Cowan I (65.0%, movie S2) and 6850 (82.4%, movie S3), PA was typically pushed aside and overgrown in interactions with JE2 (92.0% of all cases, movie S4). These results suggest that JE2 reacts more aggressively towards PA than the other two SA strains.

**Figure 4.**
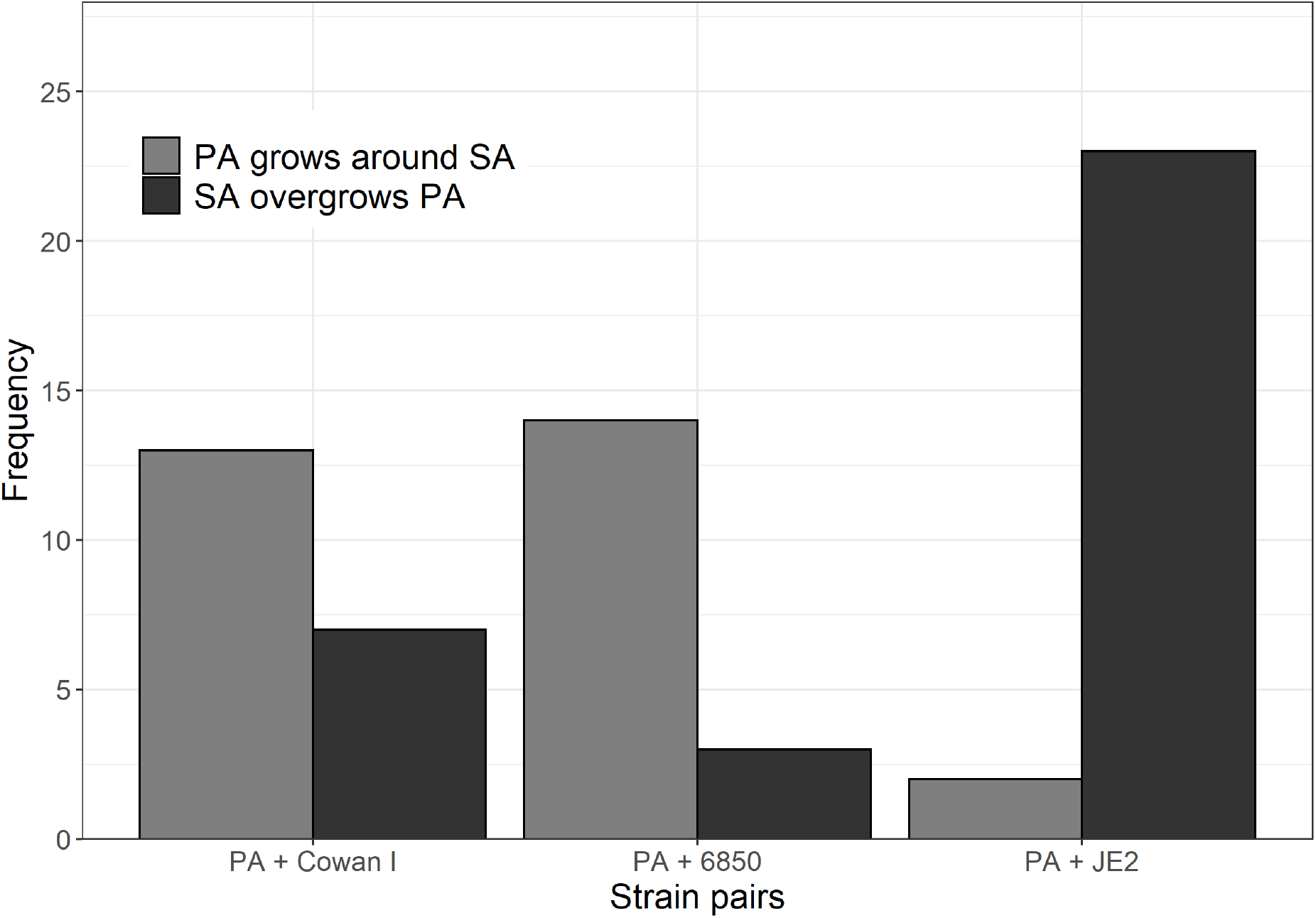
Behavioral patterns when *P. aeruginosa* (PA) and *S. aureus* (SA) microcolonies come into close contact with each other. We scanned all time-lapse image series and manually counted the frequency of the following two events: (1) PA grows around SA microcolonies (n = 29), and (2) SA pushes aside and (sometimes) overgrows PA microcolonies (n = 33). The frequency of these two types of events are significantly different across the three strain pairs (Fisher’s exact test p < 0.0001). Visual examples for the two behavioral patterns can be found in the supplementary movies 2-4 (event 1: Movie S2 and S3; event 2: Movie S4). As a comparison, Movie S1 shows PA growing in monoculture. Data is from three independent experiments per PA-SA strain pair.

### *P. aeruginosa* expedites the induction of quorum sensing systems in the presence of *S. aureus*

We hypothesized that PA might sense the presence of competitors like SA and accelerate the expression of competitive traits. To test this hypothesis, we focused on PA quorum sensing (QS) systems, which control the expression of competitive traits including the staphylolytic protease LasA and broad-spectrum toxins such as phenazines and hydrogen cyanide (44). We quantified the expression of the two main QS-regulator genes *lasR* and *rhlR* together with the housekeeping gene *rpsL* (as a control) in PA cells growing as microcolonies in the presence or absence of the three SA strains.

We found that *lasR* gene expression depended on the presence vs. absence of SA strains (ANOVA: F_3,32168_ = 697.99, p < 0.0001), the timepoint measured (5 vs. 8 hours post-inoculation: F_1,32168_ = 1394.89, p < 0.0001), and the interaction between the two (F_3,32168_ = 4621.03, p < 0.0001) (Figure 5a). Specifically, our data shows that *lasR* is induced earlier in the presence of SA (5^th^ hour), while gene expression profiles evened out later (8^th^ hour). The expression of *rhlR* was similarly affected as *lasR*. There were significant effects of the presence of SA strains (ANOVA: F_3,32331_ = 825.88, p < 0.0001), the timepoint measured (F_1,32331_ = 14818.31, p < 0.0001), and an interaction between the two (F_3,32331_ = 999.55, p < 0.0001) (Figure 5b). While *rhlR* was already expressed at the first timepoint (5^th^ hour), we also found that expression levels were generally higher in the presence of SA at both timepoints measured.

**Figure 5.**
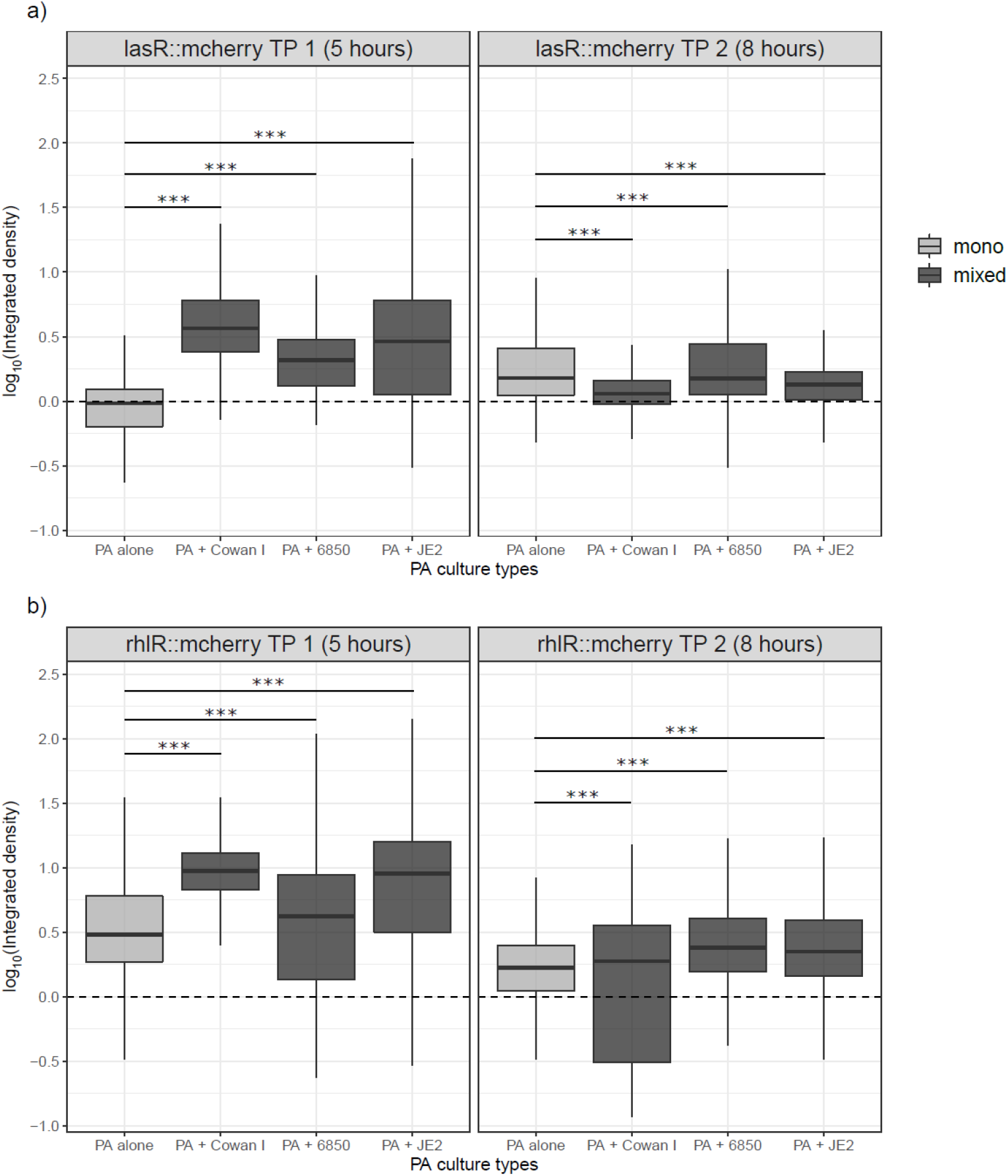
Expression of quorum sensing (QS) regulator genes in *P. aeruginosa* (PA) in mono- and mixed cultures with *S. aureus* (SA) strains. We used PA strains harboring transcriptional double reporter strains, where the genes of the QS-regulators *lasR* or *rhlR* are fused to mCherry and the housekeeping gene *rpsL* is fused to GFP (*lasR::mcherry-rpsL::gfp* and *rhlR::mcherry-rpsL::gfp*). We inoculated these strains with (dark-grey) and without (light-grey) SA strains on agarose patches and took pictures of growing microcolonies at two timepoints, after five hours (TP 1) and eight hours (TP 2) incubation at 37 °C. (a) The expression of *lasR* is increased in mixed compared to monocultures of PA after five hours, but evens out across treatments after eight hours. (b) The expression of *rhlR* is increased in mixed- compared to monocultures of PA after five hours, but evens out across treatments after eight hours. In comparison, the housekeeping gene *rpsL* is much more homogeneously expressed across all treatments and time points (Supplementary Figure S1). Supplementary Figures S2 and S3 show the expression of *lasR*, *rhlR*, and *rpsL* from both timepoints across all three independent experiments. The box plots show the median (bold line) with the first and the third quartiles. The whiskers cover the 1.5* inter-quartile range (IQR) or extend from the lowest to the highest value if they fall within the 1.5* IQR. *** p < 0.001. Data is from three independent experiments.

For both the *lasR* and the *rhlR* gene, we observed that the presence of Cowan I and JE2 had a greater influence on PA gene expression than the presence of 6850 (Figure 5). In the latter case, *lasR* and *rhlR* gene expression was more similar to the pattern shown in PA monoculture. Note that the *rpsL* housekeeping gene was constitutively expressed at both timepoints and across conditions (mono vs. mixed culture), indicating that the observed differences in QS gene expression are induced by the competitor (supplementary figure 1). In sum, the PA QS gene expression data shows that the presence of SA may lead to adjustments in PA *lasR* and *rhlR* gene expression, especially in competition with Cowan I and JE2, but to a lesser extent with 6850.

## Discussion

*Pseudomonas aeruginosa* (PA) and *Staphylococcus aureus* (SA) frequently occur together in polymicrobial infections, and there is increasing evidence that their interactions are important for virulence, disease progression, and treatment outcome (45, 46). Previous work in the field has explored molecular, ecological, and evolutionary aspects of PA-SA interactions. One key insight from this body of work is that PA is often dominant over SA through the production of a variety of inhibitory molecules (12, 13, 15, 16, 47). While these studies were mostly performed *in vitro* with planktonic batch cultures, we here used a complementary approach and studied PA- SA interactions at the single-cell level during surface-attached microcolony growth. Since during infections, PA and SA often adhere to tissues, colonize medical devices and form aggregates that develop into mature biofilms (48–50), we argue that interspecies interactions are important to study under these conditions. Using single-cell time-lapse fluorescence microscopy, we found that SA strains (Cowan I, 6850 and JE2) are highly competitive against PA. Specifically, SA cells started to divide earlier when exposed to PA and all SA strains compromised PA growth before microcolonies came into direct contact. Meanwhile, PA had little effect on SA fitness, but reacted towards the presence of SA by showing increased directional growth and increased expression of quorum sensing (QS) regulators. There were also strain- specific patterns, with PA cells growing around microcolonies of Cowan I and 6850, while being rapidly usurped by JE2 microcolonies. Altogether, our results show that on surfaces, the competitive balance tips in favor of SA (see Table 1 for a summary of all effects).

The key (and rather unexpected) finding of our study is that SA dominates PA on surfaces, which opposes the frequently observed result of PA inhibiting and outcompeting SA in planktonic batch cultures (12, 13, 15–17). One reason for why SA could be more competitive on surfaces is that this bacterial species has a non-motile lifestyle and might thus be well adapted to rapidly colonize surfaces outside and inside a host (50, 51). In contrast, PA is a flagellated motile bacterium that first engages in surface sensing to then alter its lifestyle and gene expression profile (48, 52). Surface sensing takes time and is likely associated with metabolic costs, which could put PA at a disadvantage compared to SA.

What could be the mechanisms deployed by SA to suppress PA? The fact that all SA strains started to divide earlier in the presence of PA suggests that SA can sense the presence of the competitor (53) and accelerate metabolism to trigger an earlier onset of growth (54). Although the mechanism by which SA senses competition remains to be elucidated, our observation of an earlier onset of growth indicates that SA engages in resource competition, as predicted for interactions between pathogens competing for limited host resources (55). Moreover, our observation that SA inhibits PA prior to microcolonies coming into contact suggests that SA further engages in interference competition via diffusible compounds to displace its competitor. Candidate inhibitory compounds released by SA are the phenol-soluble modulins (PSMs). PSMs are amphipathic surfactant peptides that can lyse eukaryotic and certain prokaryotic cells, they are pro-inflammatory, play a role in biofilm formation and promote SA spreading on surfaces (56, 57). PSMs are produced by virtually all SA strains, and they have previously been suggested to play a role in surface interactions with PA (38). When PA and SA cells came into contact, we saw that JE2 showed a particularly aggressive response towards PA. While we do not know whether contact-dependent interference mechanisms were involved, it was astonishing to see how PA cells were simply pushed aside and sometimes completely disappeared from the microscope field of view. It is known that community-acquired methicillin-resistant *S. aureus* (MRSA) strains, such as JE2, produce particularly high levels of PSMs (58, 59). If PSMs were indeed involved in competition on surfaces with PA, this could explain why JE2 was the most aggressive SA strain towards PA.

We now turn to PA and ask why this otherwise very competitive pathogen is comparatively weak in competition against SA on surfaces. PA features many interference traits that could harm its competitor, including LasA protease, pyocyanin, and HQNO (5). However, these interference compounds are regulated by QS and are only deployed once a certain cell density is reached (60, 61). Hence, it might be that PA is simply not ready for competition during microcolony formation. Nonetheless, PA was not idle and managed to suppress 6850, the slowest growing SA strain. This suggests that, not only SA, but also PA secretes at least some inhibitory compounds early on during competition. At later stages of microcolony formation, when coming into close contact with SA, we observed that PA reacted to the presence of Cowan I and 6850 microcolonies and started to grow around them. The observed pattern is reminiscent of the exploratory motility phenotype described by (38). While the resolution of our time-lapse movies was not high enough to follow specific cell-to-cell interactions, it did not seem that the potential exploratory motility was associated with any form of contact-dependent killing, and the benefit of this behavior thus remains to be further explored. Finally, our results indicate that PA also seems to sense its SA competitors and to mount a response through the earlier induction of QS. Interestingly, this induction was more prominent in response to Cowan I and JE2 than towards 6850, the weakest SA strain, which suggests that PA adjusts its response relative to the aggressiveness of its competitor.

We advocate the view that studying pathogen interactions on surfaces mimics more closely potential interactions in infections. However, our study is just an initial step towards a better understanding of how pathogens, such as PA and SA, might grow and interact on host-associated surfaces in infections. There are several aspects that should be considered in future studies. First, we know that relative frequencies of PA and SA impact competitive interactions in planktonic cultures (20). It would thus be important to test the effect of relative species frequencies on competition on surfaces. One possible outcome could be positive frequency- dependent competition behavior: with a high initial SA frequency, PA would probably grow very poorly, while at a high initial frequency PA might be more competitive, potentially able to keep SA at bay. Such insights could reveal so-called ‘order effects’ that are relevant for polymicrobial infections, whereby the pathogen species that colonizes the host niche first is more competitive compared to a later arriving species. Another aspect that should be investigated in more detail is the PA gene expression profile in the presence of SA. Studies on transcriptional responses of PA towards SA exist for planktonic culturing conditions, biofilms, and *in vivo* growth (15, 62, 63), and it would be important to know how the results compare to transcriptome profiles of single cells on surfaces. Interesting PA candidate genes are not only related to QS (as studied here), but also genes involved in stress response or virulence, all of which could trigger competition sensing and responses towards a competitor (53). Furthermore, little is known about secreted compounds from SA that might inhibit PA. Identifying the involvement of PSMs and possibly other SA inhibitory molecules is essential to understand how SA suppresses PA on surfaces. Finally, while we looked at the early stages of microcolony formation, it would be interesting to look at strain dynamics in more mature biofilms, for instance by using flow chambers combined with confocal microscopy 3D analysis, where experiments can be run for longer without the cells overgrowing each other, which frequently occurs after prolonged hours of microcolony growth using our agarose patches.

Taken together, our work shows that the two human opportunistic pathogens *P. aeruginosa* (PA) and *S. aureus* (SA) influence each other at the single-cell level on surfaces in manifold ways. While both species seem to be able to sense competition, SA was more competitive, showing both signs of resource competition by starting to grow earlier, and interference competition through diffusible compounds reducing the growth of PA. Crucially, SA is much more competitive on surfaces than would be anticipated from planktonic batch culture experiments. Since PA and SA colonize host tissues in the context of infection, we provide new hints on the competitive strengths of these two important pathogens that often co-exist in infections. Moreover, our results with a panel of genetically distinct SA strains suggests that the virulence potential of SA strains might play a role in competition with PA, with JE2 being the most virulent and most competitive strain on surfaces. Overall, we propose that time-resolved quantitative live imaging has the potential to uncover novel interspecies interactions in an ecologically relevant context. This approach may not only be useful to further our insights on interactions between PA and SA but may significantly improve our understanding of interactions between any two or more species infecting a host based on a surface-colonizing lifestyle.

## Supporting information

Statistics_data

## Author Contributions

S.N. and R.K. designed research, S.N. performed research, T.W. wrote the image analysis scripts, S.N. and R.K. analyzed data and wrote the paper with input from T.W.. All authors approved the manuscript.

## Acknowledgements

We thank Markus Huemer (University Hospital of Zürich) for providing *S. aureus* strains, members of the Kümmerli Group for providing engineered *P. aeruginosa* gene reporter strains, and the Center for Microscopy and Image Analysis of the University of Zürich for technical support and maintenance of resources. Illustration for Figure 1 was created using BioRender (www.biorender.com).

## Data availability statement

All raw data sets will be deposited in the figshare repository (DOI will be provided upon the acceptance of the manuscript).

