## Supplementary_Material for "Single-cell imaging reveals that *Staphylococcus aureus* is highly competitive against *Pseudomonas aeruginosa* on surfaces"

**Supplementary Table 1.** Strains used for this study.

| Species and strain name | Origin | Description | Fluorophore | Reference |
|---|---|---|---|---|
| ***Pseudomonas aeruginosa* (PA)** | | | | |
| PAO1 | Wound | Commonly used *P. aeruginosa* laboratory strain. | (1) *ptac::gfp*<br>(2) *Promoterless::mcherry-Promoterless::gfp*<br>(3) *lasR::mcherry-rpsL::gfp*<br>(4) *rhlR::mcherry-rpsL::gfp* | ATCC 15692 |
| ***Staphylococcus aureus* (SA)** | | | | |
| Cowan I | Septic arthritis | MSSA isolate. Highly invasive, but not cytotoxic. Agr-defective. | untagged | ATCC 12598 |
| 6850 | Osteomyelitis | MSSA isolate. Highly invasive, cytotoxic, and hemolytic. | untagged | ATCC 53657 |
| JE2 | Skin and soft tissue infection | USA300 CA-MRSA isolate. Highly virulent, cytotoxic, and hemolytic. | untagged | NARSA |

All fluorescent constructs in PAO1 were chromosomally inserted using the mini-Tn7 insertion system.

CA-MRSA: Community-acquired methicillin-resistant *S. aureus*

MSSA: Methicillin-sensitive *S. aureus*

Agr: Accessory gene regulator

Supplementary Figure 1

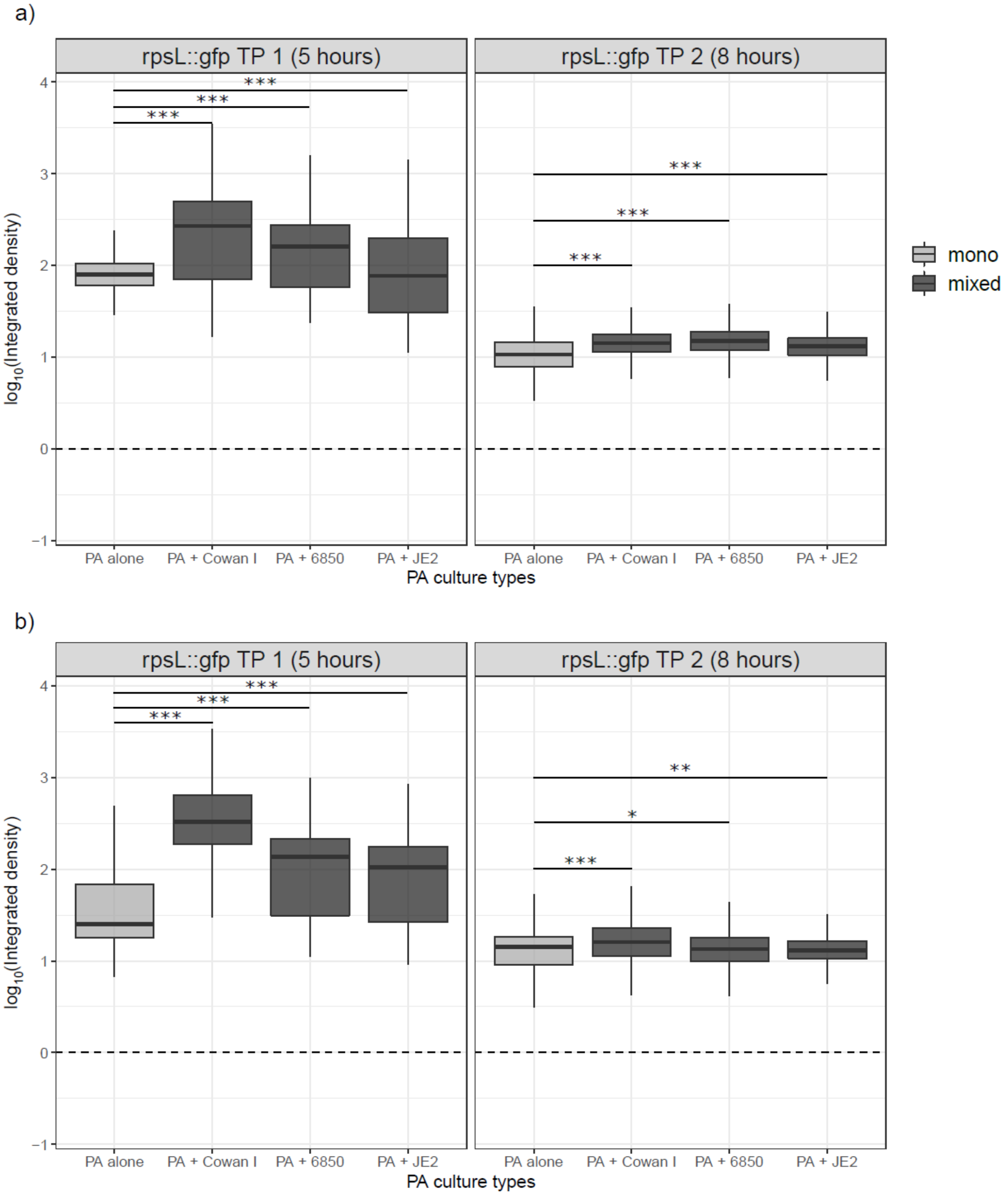


**Supplementary Figure 1.** Expression of *rpsL::gfp* in the PA *lasR* and *rhlR* double reporters with and without SA. We used PA strains harboring transcriptional double reporter strains, where the genes of the quorum sensing regulators *lasR* or *rhlR* are fused to mCherry and the housekeeping gene *rpsL* is fused to GFP (*lasR::mcherry-rpsL::gfp* and *rhlR::mcherry-rpsL::gfp*). We inoculated these strains with (dark-grey)

and without (light-grey) SA strains on agarose patches and took pictures of growing microcolonies at two timepoints, after five hours (TP 1) and eight hours (TP 2) incubation at 37 °C. a) Expression of *rpsL* in the *lasR::mcherry-rpsL::gfp* reporter is more homogeneous than expression of *lasR*. b) Expression of *rpsL* in the *rhlR::mcherry-rpsL::gfp* reporter is more homogeneous than expression of *rhlR*. The box plots show the median (bold line) with the first and the third quartiles. The whiskers cover the 1.5* inter-quartile range (IQR) or extend from the lowest to the highest value if they fall within the 1.5* IQR. *** $p < 0.001$, ** $p < 0.01$, * $p < 0.05$. Data is from three independent experiments.

**Supplementary Figure 2**

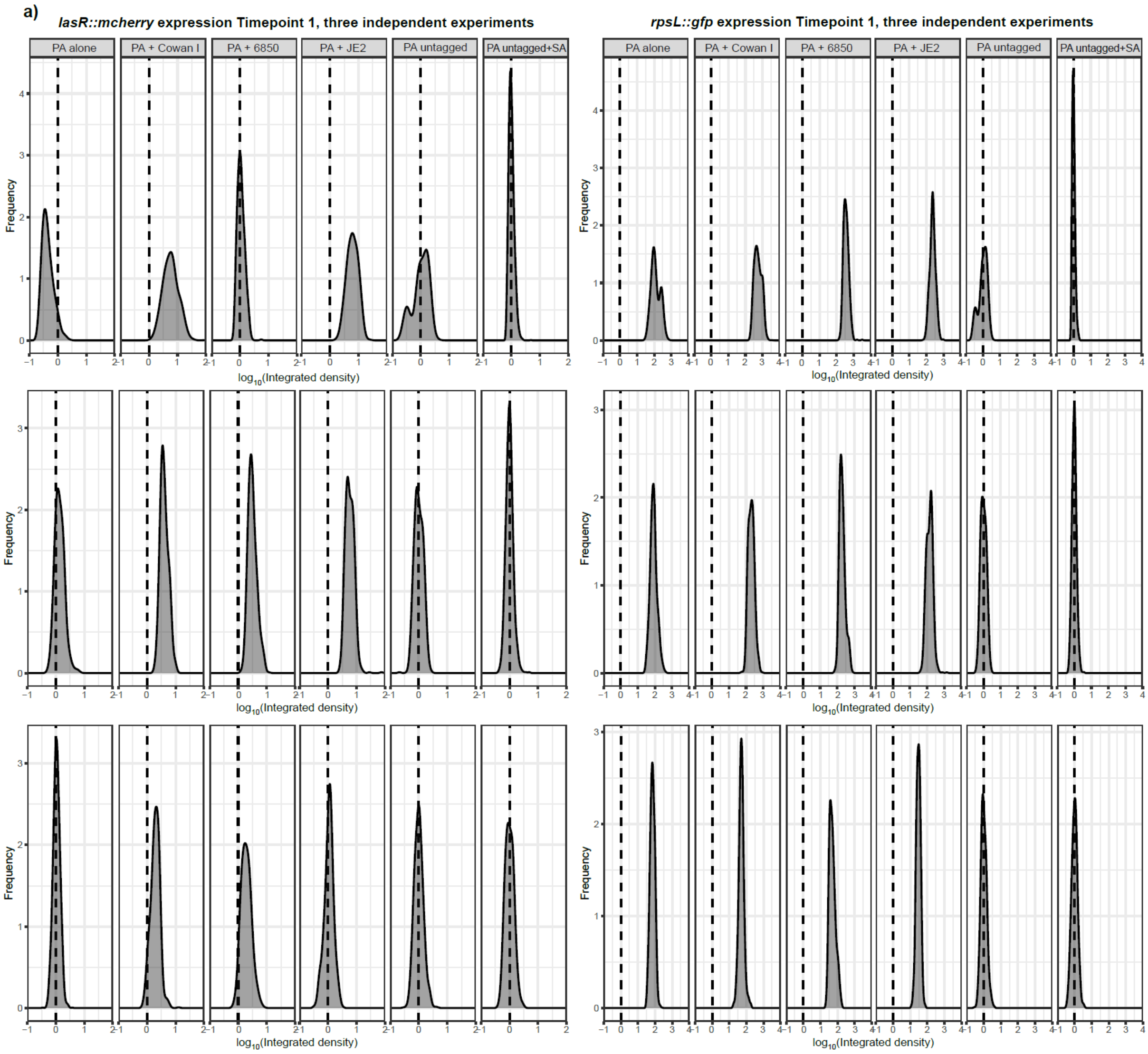

b)

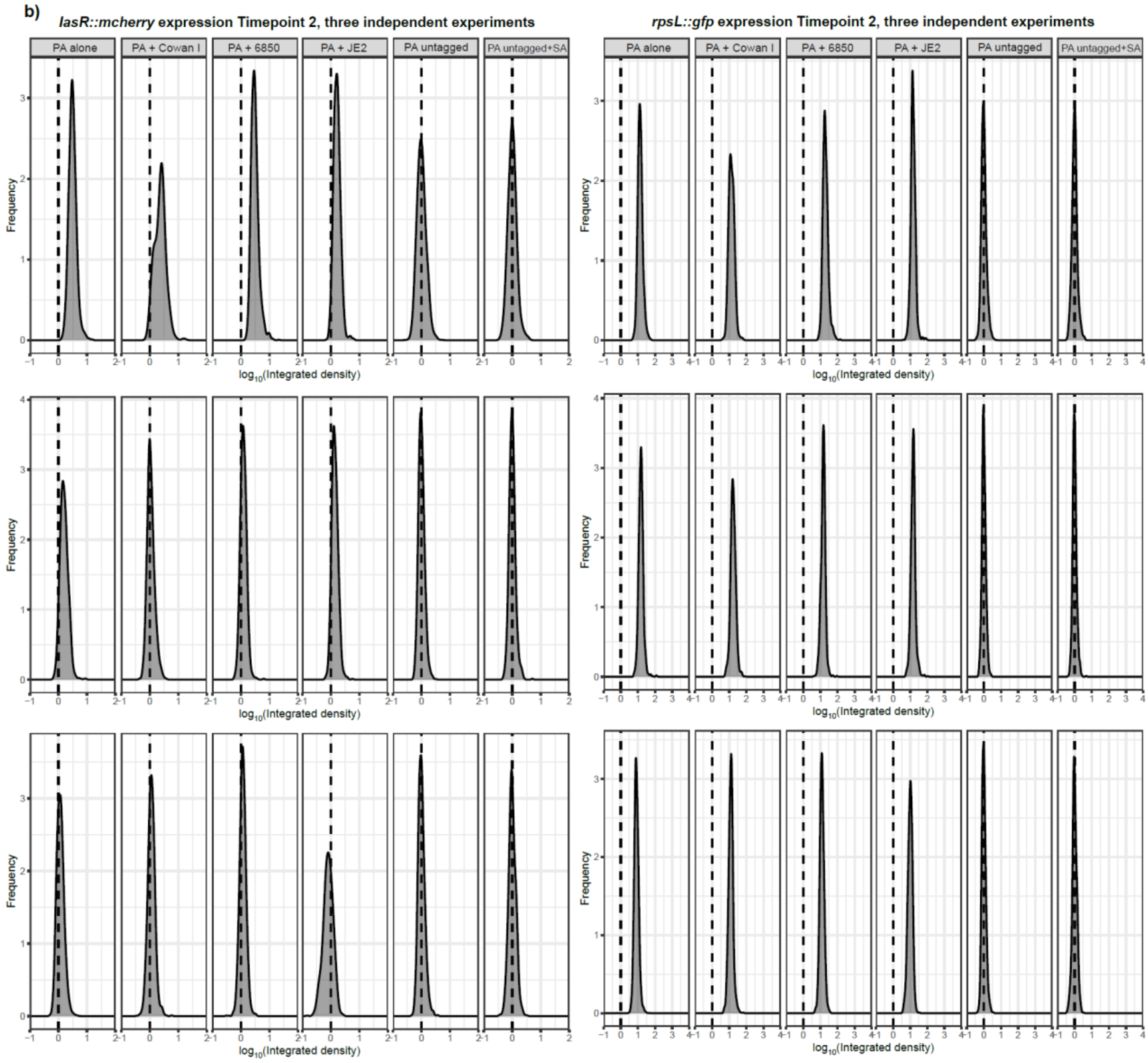


**Supplementary Figure 2.** Individual histograms for *lasR::mcherry* (left) and *rpsL::gfp* (right) expression in the PA double reporter strain *lasR::mcherry-rpsL::gfp.* a) Timepoint 1 (five hours); b) Timepoint 2 (eight hours). Data is from three independent experiments (each row corresponds to one independent experiment).

**Supplementary Figure 3**

**a)**

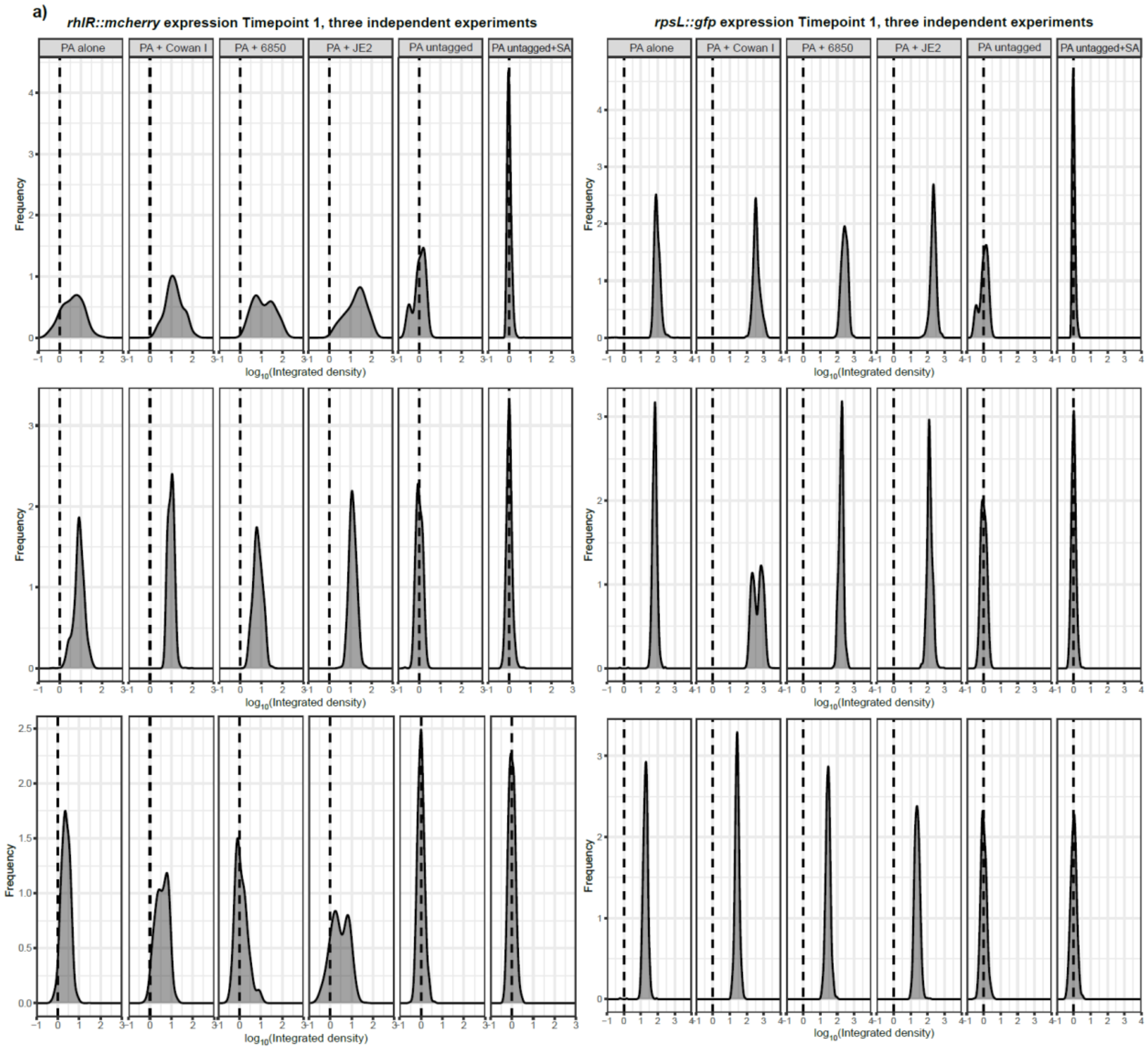

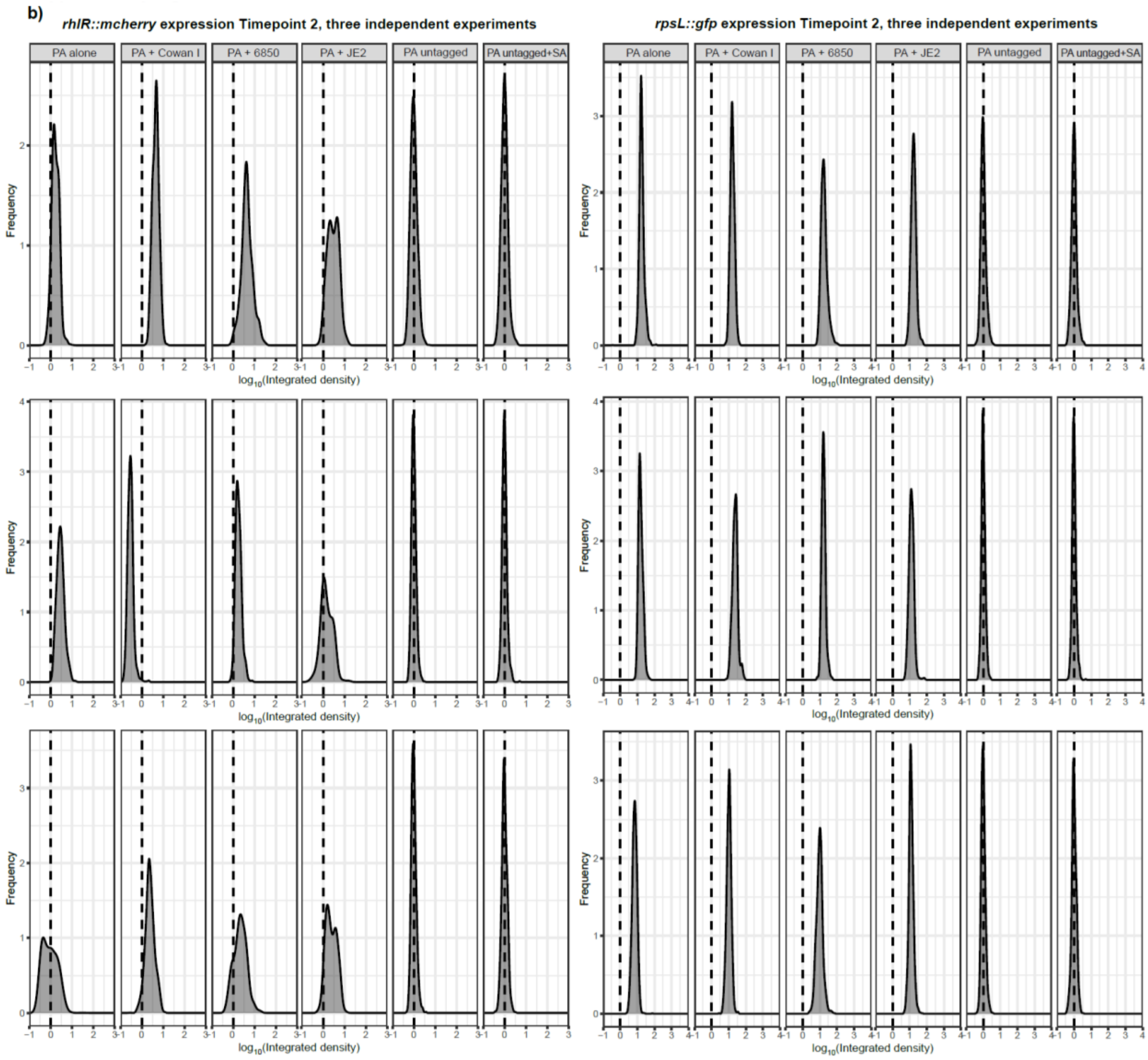


**Supplementary Figure 3.** Individual histograms for *rhlR::mcherry* (left) and *rpsL::gfp* (right) expression in the PA double reporter strain *rhlR::mcherry-rpsL::gfp.* a) Timepoint 1 (five hours); b) Timepoint 2 (eight hours). Data is from three independent experiments (each row corresponds to one independent experiment).

**Supplementary Movies 1-4.**

All time-lapse movies were acquired over six hours at 37 °C with pictures taken every 10 min. Scale bar at the lower right of each movie represents 10 µm. For clarity, only the phase contrast channel is shown.

**Movie S1.** *P. aeruginosa* (PA) growing alone.

**Movie S2.** PA together with the *S. aureus* (SA) strain Cowan I.

**Movie S3.** PA together with the SA strain 6850.

**Movie S4.** PA together with the SA strain JE2.
